# *Syn3* Gene Knockout Negatively Impacts Aspects of Reversal Learning Performance

**DOI:** 10.1101/2021.05.21.445145

**Authors:** Alyssa Moore, Jérôme Linden, James D. Jentsch

## Abstract

Behavioral flexibility enables the ability to adaptively respond to changes in contingency requirements to maintain access to desired outcomes, and deficits in behavioral flexibility have been documented in many psychiatric disorders. Previous research has shown a correlation between behavioral flexibility measured in a reversal learning test and *Syn3*, the gene encoding synapsin III, which negatively regulates phasic dopamine release. *Syn3* expression in the hippocampus, striatum, and neocortex is reported to be negatively correlated with reversal learning performance, so here, we utilized a global knockout line to investigate reversal learning in mice homozygous wildtype, heterozygous null, and homozygous null for the *Syn3* gene. Compared to wildtype animals, we found a reversal specific effect of genetic *Syn3* deficiency that resulted in a greater proportional increase in trials required to reach a preset performance criteria during contingency reversal, despite no observed genotype effects on the ability to acquire the initial discrimination. Behavioral flexibility scores, which quantified the likelihood of switching subsequent choice behavior following positive or negative feedback, became significantly more negative in reversal only for *Syn3* homozygous null mice, suggesting a substantial increase in perseverative behavior in the reversal phase. *Syn3* ablation reduced the number of anticipatory responses made per trial, often interpreted as a measure of waiting impulsivity. Overall, *Syn3* expression negatively affected behavioral flexibility in a reversal specific manner but may have reduced waiting impulsivity.

## Introduction

Behavioral flexibility relates to an individual’s ability to modify behavioral patterns to fit changing environmental conditions. Deficits in behavioral flexibility have been characterized in a number of psychiatric disorders (Uddin, 2021), including schizophrenia (Waltz, 2017; Waltz & Gold, 2007), autism spectrum disorder (Kelly & Reed, 2020), obsessive-compulsive disorder (Gruner & Pittenger, 2017; Vaghi et al., 2017), and substance use disorders (Winstanley et al., 2010; Isten et al., 2017; Izquierdo & Jentsch, 2012; Robbins et al., 2012; Ersche et al., 2010).

Reversal learning is an operant test of behavioral flexibility in which an initial association (stimulus-response or response-outcome) is learned through reinforcement, before the conditions for reinforcement are reversed and the organism is tested for its ability to update behavior (Izquierdo et al., 2017, Izquierdo & Jentsch, 2012). In other words, one response is established as prepotent during initial acquisition through positive feedback, and this action must then be inhibited or changed during reversal testing to procure reward.

Dopaminergic tone has been implicated in the ability to reverse a learned discrimination (Klanker et al., 2013). Klanker et al. (2015) identified a key role for ventromedial striatal dopamine (DA) responses to positive feedback while animals acquired a spatial reversal task. In rats that eventually acquired the reversal rule, the first rewarded reversal trial induced a spike in DA (measured via fast-scan cyclic voltammetry) concurrent with reward delivery; in the subsequent trial (i.e., the first trial in which positive feedback can be used to update behavior), an increase in cue-evoked DA was observed. In rats that did not acquire reversal, the first reward induced the same DA spike, but DA release did not shift to cue presentation in the next trial. This effect was specific to positive feedback; DA release during an incorrect trial (i.e., one followed by no reward delivery) or the following trial did not differ between rats that acquired the reversal and those that did not. A change point was defined as the trial at which the cumulative record maximally deviated from a line drawn from the origin to the end of the record (Klanker et al., 2015). No differences were observed in cue-evoked DA release in trials before and after the change point, but reward-evoked DA release decreased in trials following the change point. Positive feedback increased cue-evoked DA release on the subsequent trial only prior to the change point, suggesting feedback-induced shifts in intra-trial timing normalize across the learning curve to a level where cue-evoked DA stabilizes, and the learned behavior is reliably expressed. Dopaminergic tone is dynamically engaged throughout the learning curve, and small shifts in the timing of release can aid in reorganization of previously learned behavior in response to positive feedback.

Yawata et al. (2012) demonstrated independent roles for the direct (D1 receptor-expressing) and indirect (D2 receptor-expressing) dopamine-mediated pathways of the basal ganglia by reversibly inducing pathway specific blockade of neurotransmission in the nucleus accumbens using doxycycline in transgenic mice. Interference with the D1 receptor-expressing direct pathway impaired acquisition of a novel visual discrimination, though no impairment in acquisition was observed when the D2 receptor-expressing indirect pathway was blocked. Blocking neurotransmission in either the direct or indirect pathway interfered with reversal of the previously learned discrimination, but only inhibiting the indirect pathway neurons increased perseverative errors. This pattern of findings was recapitulated when the direct or indirect pathways were unilaterally blocked via doxycycline, then the contralateral side treated with D1 or D2 agonists or antagonists. Antagonism of accumbal D1 receptors interfered with the acquisition of initial and reversed discriminations but did not alter perseverative errors, while D2 agonism selectively hindered reversal performance by increasing perseverative errors (Yawata et al., 2012). Together, these results suggest the D1-mediated direct pathway in the basal ganglia is involved in the acquisition of operant responses more generally, while DA release onto the D2-mediated indirect pathway is specifically implicated in behavioral flexibility.

Lee et al. (2007) demonstrated that in monkeys systemic antagonism of D2/D3, but not D1/D5, specifically impaired performance following reversal of a previous learned visual discrimination without altering the ability to acquire novel discriminations. In rats, intra-accumbal D1, but not D2, antagonism disrupted set-shifting (a measure of cognitive flexibility), while D2, but not D1, agonism impaired performance on a reversal task without disrupting initial discrimination (Haulk & Floresco, 2009). Although one might expect agonism and antagonism of D2 receptors to have opposite effects, conceptual and experimental evidence points to a convergent role of the *change* in dopamine activity to drive behavioral effects. Because D2 receptors are found both pre- and post-synaptically, agonism will inhibit further dopamine release via autoreceptors, while antagonism will interfere with post-synaptic activation even as presynaptic release is disinhibited. In both cases, phasic dopamine activity is disrupted, and reversal learning is impaired.

### Genetic correlates of reversal learning

Laughlin et al. (2011) evaluated reversal learning performance in a panel of BXD mouse strains and utilized a genome-wide linkage approach to model the impact of genetic variation on the reversal phenotype. A genome-wide quantitative trait locus on mouse chromosome 10 was identified, and *Syn3* emerged as a provocative positional candidate expressed from that genomic region. *Syn3* mRNA expression is regulated in *cis*, and its expression in the hippocampus, neocortex, and striatum was found to be positively correlated with reversal learning performance in the BXD panel, such that greater *Syn3* expression correlated with faster reversal learning (Laughlin et al., 2011). Synapsin III, encoded by the *Syn3* gene, is a member of the synapsin family of neuronal phosphoproteins (Kao et al., 1998). Synapsin III can be localized on the cytoplasmic side of synaptic vesicles and is implicated in neurotransmitter release. Feng and colleagues (2002) demonstrated that loss of synapsin III led to larger vesicular recycling pools but did not alter vesicular release or quantal dynamics. This view is supported by concurrent evidence that synapsin proteins regulate a distal reserve pool of vesicles (Greengard et al., 1993; Hilfiker et al., 1999; Hilfiker et al., 2005). Functionally, a loss of synapsin III prevents vesicles from being sequestered away from the ready-release pool, promoting more sustained release in response to continued stimulation. In a typical case, sustained release would be limited by the rate of transfer of vesicles from the reserve pool to the active zone; this rate-limiting process is theoretically disrupted in cells lacking synapsin III as vesicles are inadequately sequestered in the reserve.

Synapsins have been found to be differentially expressed in neuronal populations. Bogen and colleagues (2006) found that selective deletion of synapsin I and II substantially reduced vesicular uptake of GABA and glutamate but did not alter DA uptake. The differences in vesicular uptake were driven by reduced concentration of vesicular transporters related to glutamate and GABA, but no change in VMAT2, the transporter responsible for packaging dopamine into synaptic vesicles. Further, synapsin I and II were found to colocalize in cells expressing vesicular transporters for GABA and glutamate, but dopaminergic terminals appeared not to colocalize with synapsin I and II. Synaptosomal uptake was enhanced in all three transmitter systems in synapsin I and II knockouts compared to wildtype controls, adding additional complexity to the role of synapsins in neurotransmission (Bogen et al., 2006). A subsequent study utilized a triple knock-out approach to investigate differential regulation of dopamine and serotonin by synapsins (Kile et al., 2010): serotonin was not altered by deletion of all three synapsins, but DA release was significantly enhanced. Selective deletion of synapsin III also elicited enhanced DA release, demonstrating a distinct role for the synapsins in regulating neurotransmitter release. Because DA is extensively implicated in neuropsychiatric disorders, subcellular proteins which contribute to dopamine dynamics and the genes encoding them are of intense interest.

### Hypothesis

In this study, we investigated the role of *Syn3* on reversal learning performance using a genetic knockout strategy. Mice expressing 0, 1, or 2 functional *Syn3* alleles were tested for reversal learning. We hypothesized *Syn3* function would be related to behavioral flexibility, such that mice lacking functional *Syn3* would exhibit deficits in reversal learning performance without showing impairments on the acquisition of the initial discrimination.

## Methods

### Animals

C57BL/6N-Syn3^tm1.1(KOMP)Vlcg^/J mice (RRID:MMRRC_049950-UCD) were obtained from The Jackson Laboratory (Bar Harbor, ME). They were maintained through heterozygote crosses, producing all 3 genotypes (homozygous null, heterozygous, or homozygous wildtype, having 0, 1, or 2 functional *Syn3* alleles, respectively) in each litter. They were housed in a temperature and humidity-controlled vivarium on a 12-hour light-dark cycle, with all procedures being conducted in the light phase.

Mice were weaned on postnatal day 21, at which point they were housed in same-sex groups of 3-5 mice per cage. Offspring were genotyped for dosage of wildtype and mutant *Syn3* alleles by Transnetyx (Cordova TN) from ear tissue collected at weaning.

A total of 112 mice offspring were involved in the study, but 6 were removed because they failed to meet pre-set discrimination performance criteria or due to experimenter errors. Mice that did not reach acquire the initial discrimination within 30 sessions were excluded from analysis (n=2, both heterozygous null). Four mice were excluded due to experimenter error. The final number of mice included in the forthcoming analyses was 106 (54 females, 52 males; 37 homozygous null, 42 heterozygous and 27 wildtype mice). All protocols were reviewed and approved by Binghamton University’s Institutional Animal Care and Use Committee, and all procedures were consistent with the <u>Guide for the Care and Use of Laboratory Animals</u> (National Research Council, 2011).

### Reversal learning

Prior to the onset of these studies, mice were briefly handled daily to be weighed and tail marked. Subsequently, mice were food restricted to ~85% of their free feeding body weights; mice were fed once per day, after testing, in an amount that was individually titrated to achieve this reduced weight. Before the start of the experiment, mice were offered ~0.5 g of reinforcer pellets (14-mg Dustless Precision pellets; stock number F05684; Bio-Serv Inc., Flemington NJ) per mouse in their home cage.

The equipment and procedures used for testing followed the protocol originally described in Laughlin et al. (2011). All testing took place in operant conditioning chambers (model MED-NP5M-D1; Med Associates; St Albans VT), each enclosed in a sound-attenuating cubicle. The chambers were equipped with a house light and white noise generator, located outside of the chamber but within the cubicle. A photocell-equipped food delivery magazine connected to a pellet dispenser were on one wall of the chamber and a horizontal array of 5 nose-poke apertures were on the opposite wall. Mice were first exposed to the testing chamber in a single 30-min habituation session, with the house light and white noise on. Magazine training began the following day. Reinforcer pellets were delivered into the internally illuminated magazine at the start of these sessions, and again every 30 s after each pellet was retrieved, until 50 pellets were delivered or 45 min passed, whichever occurred first. Mice remained in magazine training until they retrieved 50 pellets in a single session.

A three-stage aperture training followed. In the first stage, the food magazine was illuminated at the start of the session, and head entry resulted in pellet delivery, termination of magazine illumination, and illumination of the central nose poke aperture on the opposite wall. A response into the lit central nose poke terminated the nose poke illumination and led to illumination of the magazine and delivery of a pellet; a variable observing response (OR) nose poke time of 0, 10, 20, or 40 cs was required, randomized from trial to trial. Stages 2 and 3 followed the same general schedule, progressively increasing the OR duration array. In Stage 2, the observing response array increased as a function of the rewards earned. When fewer than 15 reinforcers had been earned, an OR of 0, 10, 20, or 40 cs could be required. For reinforcers 16-25, OR times could be 0, 20, 30, or 50 cs. For the remainder of the session, OR durations could be 0, 20, 40, or 60 cs. Stage 3 OR durations were 20, 40, or 60 cs throughout the session. Transition from one stage to the next required mice to earn at least 30 reinforcers in a single session.

Mice began discrimination acquisition training the day following completion of stage 3. Here, an observing response in the central nose poke aperture (variable hold requirement of 20, 40, or 60 cs) initiated a trial, at which point the aperture holes flanking the center were both illuminated. Mice had 30 seconds to make an entry into one of those two apertures and were reinforced with two pellets for selecting the “correct” one and punished with a 5-s time-out with all visual stimuli off for selecting the “incorrect” one. Which of the two apertures was selected to be “correct” was pseudo-randomly assigned for each mouse and maintained throughout discrimination training. Failure to respond within the 30-s window was scored as an omission. Omissions were followed by a 5-s time-out. Correct and incorrect responses, as well as omissions, were followed by a 3-s inter trial interval (ITI), during which no apertures were illuminated. Each session lasted for 125 trials, 60 min, or until discrimination criteria was met, whichever occurred first. The criterion for completing discrimination acquisition training was 80% correct responses in a sliding window of 20 trials. Mice that made fewer than 5 responses during two consecutive days of discrimination acquisition training were returned to Stage 2 of aperture training, and then returned to the discrimination phase after passing aperture training again. Mice failing to respond in the reversal phase were not transferred back to training but were removed from the study. This did not apply to any mice tested in this experiment.

Reversal began the day after mice completed discrimination training. Testing conditions were identical, except that the aperture that resulted in pellet delivery during discrimination no longer produced reward and the opposite aperture now resulted in delivery of two pellets. Animals were tested until reaching the same performance criterion used during acquisition.

### Dependent variables

A series of calculated dependent variables are used to evaluate the performance of individual mice in the test; all variables are calculated separately for the acquisition and reversal stages. The number of trials required to reach the preset performance criterion is a key dependent variable, as this was the variable subject to the genome wide linkage studies in Laughlin et al. (2011) that led to the selection of *Syn3* as a positional candidate gene. Higher trait values are indicative of more difficulty with learning the initial or reversed rule.

To better understand the impact of contingency reversal on the number of trials required to reach performance criteria, a reversal fold-change variable was calculated as the number of trials to criterion in the reversal phase divided by the number of trials in the initial discrimination phase. This variable communicates the same information as a traditional difference score, but cannot include negative numbers, making it more amenable to transformation. A negative value in a difference score would indicate fewer trials were needed to reach criteria in reversal than in discrimination; a reversal fold-change score less than 1 indicates the same response pattern.

To investigate the influence of prior outcome on future choices, a behavioral flexibility score was calculated for each subject (Aarde et al., 2019). The behavioral flexibility score quantifies the tradeoff between flexibility and stability in choice behavior. The “correct” response following a reward delivery is to make the same choice again (“Win-Stay” or success through stability), while the correct choice following a timeout is to make the opposite choice on the next trial (“Lose-Shift” or success through flexibility). We evaluated flexibility in response to positive and negative feedback by determining the relative likelihood of shifting behavior on the subsequent trial. Behavioral flexibility scores were calculated independently for positive and negative feedback as (shift trials – stay trials) / (shift trials + stay trials), and are bound by −1 and 1, with −1 indicating that subject never shifted responding, and 1 indicating they always shift. In our task, the optimal flexibility score following positive feedback is −1; and optimal flexibility following negative feedback is +1.

Anticipatory responses are responses made to the correct or incorrect apertures after one trial has been completed but before the next one has been started by a satisfactory OR (i.e., during the ITI or trial initiation periods). These responses were counted and expressed as a fraction of the total number of trials initiated. As noted above, OR of minimum duration were required to initiate a trial; in some cases, mice broke their OR before reaching the minimum criterion, and these OR failures were counted as a fraction of the number of presentations of each observing response duration.

A number of latency measures were collected to allow for deeper analysis of reversal learning performance and progression through stages of the task. Trial initiation latencies were measured as the time (in ds) from the start of a new trial to initiation of the observing response. Response latencies were measured as the time (in cs) between the presentation of the target apertures and the selection of one option; these were divided into correct and incorrect latencies, based upon the outcome of the trial. Pellet retrieval latencies were measured as the time (in ds) between pellet delivery following a correct response and head entry into the magazine.

### Statistical analyses

Generalized estimating equations (GEE) were used to model the effect of genotype, sex, phase, and their interactions on performance. The model used the robust estimator for the covariance matrix, an unstructured correlation matrix, the maximum likelihood method of parameter estimation, Type III model effects, and the Chi-square Wald statistic for the full log quasi-likelihood function. Normality and linearity were assessed by Kolmogorov-Smirnov test and P-P plots, respectively. In cases where normality and linearity were validated, a normal distribution with identity link function was used. In cases where linearity or normality were violated, transformations were conducted and gamma and inverse gaussian distributions were tested. Intercept-only models (i.e., models lacking sex and genotype as predictors) were built for each variable to assess goodness of fit using the Corrected Quasi Likelihood under Independence Model Criterion (QICC). Table 1 shows the relevant transformations and model information for each test. Phase was not included as a within-subject predictor in reversal fold change because this variable denotes a ratio between acquisition and reversal, so phase is inherently accounted for. Pairwise *post hoc tests* were used to evaluate significant model effects on all main and interaction effects, except main effect of genotype; genotype was analyzed using a Simple contrast *post hoc* analysis with homozygous wildtype mice as the reference group. All *post hoc* tests used the Sidak correction for multiple comparisons. Means in the text are estimated marginal means (EMMs) ± SEM unless otherwise stated. Data are presented in figures as raw ± SEM unless otherwise stated.

**Table 1.** GEE model parameters.

| Dependent variable | Transformation | Probability distribution | Link function | Within-subject effect | Prediction factor |
| --- | --- | --- | --- | --- | --- |
| Trials to criterion | None | Inverse Gaussian | Identity | Phase | Phase, genotype, sex |
| Reversal fold change | None | Inverse Gaussian | Identity | -- | Genotype, sex |
| Anticipatory responses per trial | +1 (added to trials), log | Inverse Gaussian | Identity | Phase, side | Phase, side, genotype, sex |
| Omissions per trial | None | Normal | Identity | Phase | Phase, genotype, sex |
| Trial initiation latency | None | Inverse Gaussian | Identity | Phase | Phase, genotype, sex |
| Pellet retrieval latency | Box-Cox | Inverse Gaussian | Identity | Phase | Phase, genotype, sex |
| Response latency | Box-Cox | Inverse Gaussian | Identity | Phase, side | Phase, side, genotype, sex |
| Observing response failures per trial | +1 (added to observing response failures), log | Inverse Gaussian | Identity | Phase, observing response requirement | Phase, observing response requirement, genotype, sex |
| Flexibility score | +2 (added to score), log | Normal | Identity | Phase, feedback valence | Phase, feedback valence, genotype, sex |

Estimation statistics (Ho et al., 2019) were used to further explore the data. Analyses were conducted using the DABEST package in R, and Cumming estimation plots were generated using the raw data, presented as individual dots in swarm plots on each chart. Vertical lines next to the swarm plots represent mean ± SD for that group. Unpaired mean difference plots display the mean difference distributions ± SD on the y-axis, and the groups being compared on the x-axis.

## Results

### Trials to criterion

As expected, mice required significantly larger numbers of trials to reach a performance criterion in the reversal *vs*. acquisition stage [χ^2^_(1, N=106)_ = 79.248, *p* < .001; Table 2]. Our model identified no significant main effect of Genotype [χ^2^_(2, N=106)_ = 4.918, *p* = .086], nor any significant Genotype*Phase interaction [χ^2^_(2, N=106)_ = 4.957, *p* = .086], though we did identify a significant Genotype*Sex interaction [χ^2^_(2, N=106)_ = 5.997, *p* = .050]. There was no significant main effect of sex, nor any higher-level interactions involving sex (all χ^2^ < 4.957, all p > .05).

**Table 2.**
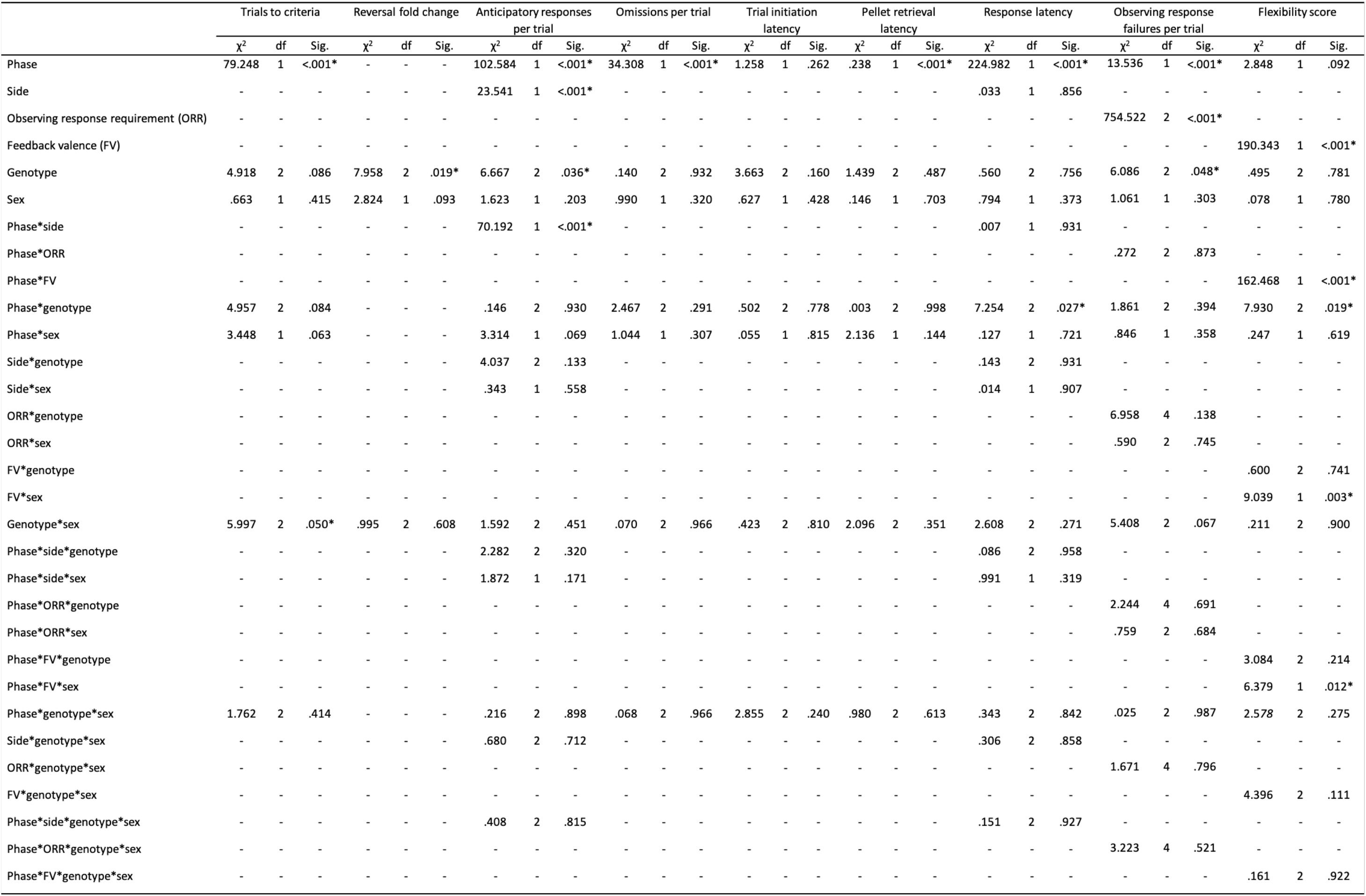
Outcomes of GEE models on multiple task measures.

An estimation statistics approach was used to further explore the data, in light of our *a priori* hypothesis. All genotypes exhibited equivalent acquisition of the initial discrimination (Fig. 1a), and all experienced more difficulty with reaching performance criteria in the reversal, as compared to initial phase (Fig. 1b, bottom panel). However, the magnitude of increase in trials to criteria does not appear to be the same across genotypes. Specifically, heterozygous null mice experienced considerable difficulty in meeting criteria following contingency reversal. To better visualize the reversal specific impairments, a fold change statistic was calculated and visualized (Fig. 1c), and a GEE model was fit to the data.

**Figure 1.**
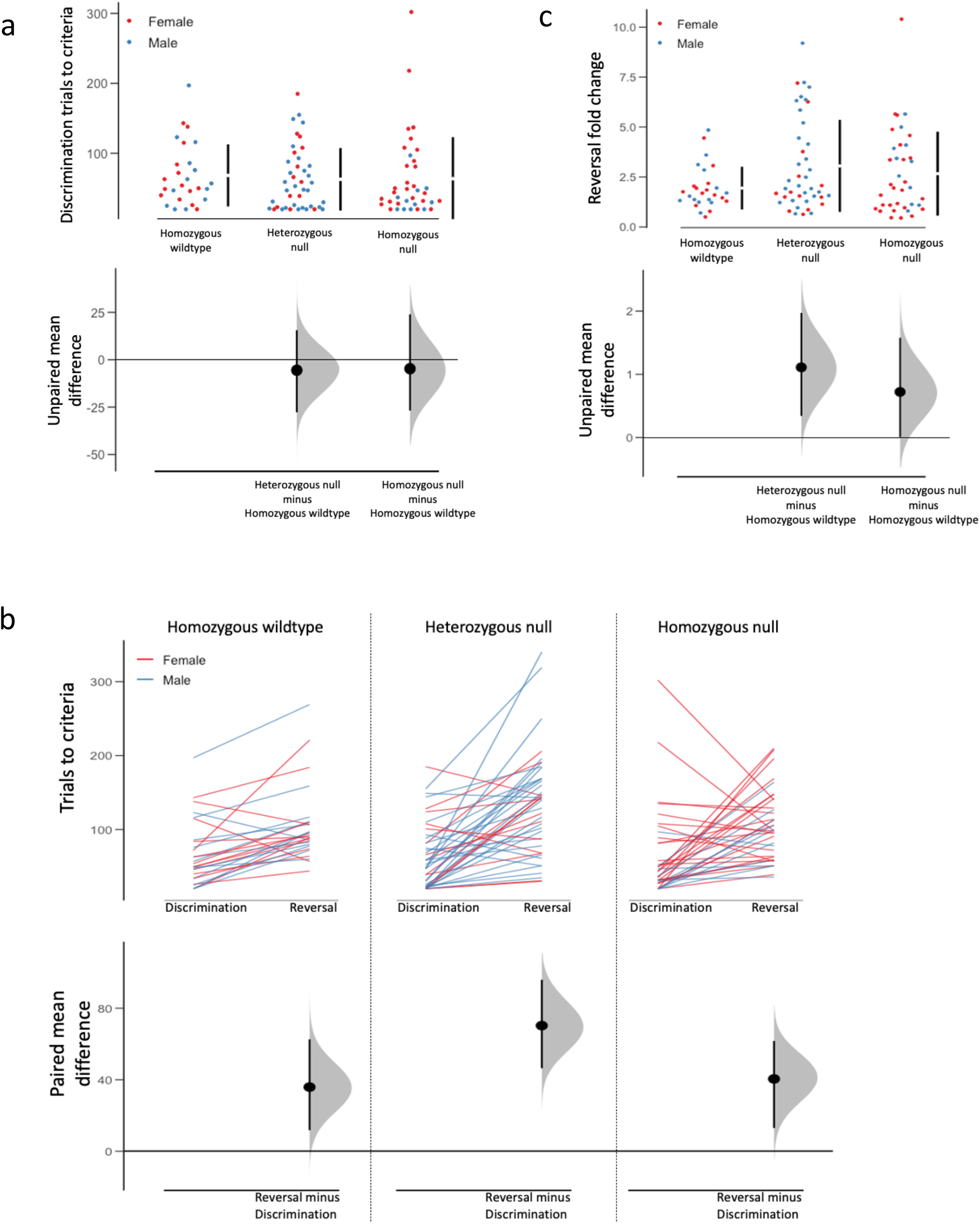
*Syn3* ablation negatively affects reversal learning performance. **(a)** Cumming estimation plot displaying discrimination performance between genotypes. The swarm plot in the top panel shows individual data points for each group with mean (±SD) displayed in the bar to the right. Unpaired mean differences comparing each mutant group to the wildtype control group are shown in the bottom panel. **(b)** Cumming estimation plot displaying discrimination and reversal performance within each genotype group. The line plot in the top panel shows individual data for mice in each genotype group. Paired mean differences comparing discrimination and reversal performance within each group are in the bottom panel. **(c)** Cumming estimation plot displaying reversal fold change score between genotype groups. The swarm plot in the top panel shows individual data points for each group with mean (±SD) displayed in the bar to the right. Unpaired mean differences comparing discrimination and reversal performance within each group.

Genotype was found to significantly influence reversal fold change [χ^2^_(2, N=106)_ = 7.958, *p* = .019; Table 2], and *post hoc* analysis revealed that reversal more substantially impaired performance in heterozygous mutant mice (mean = 2.89, SE = .37) compared to homozygous wildtype mice (mean = 1.95, SE = .26; *post hoc p* = .037). The *post hoc* contrast comparing homozygous wildtype and homozygous mutant genotypes also trended on significance (*p* = .064), and the Cumming estimation plot (Fig. 1c) shows only a very small portion of the mean difference distribution would support the null hypothesis that homozygous mutants are phenotypically equivalent to homozygous wildtype mice.

### Anticipatory responses

Results of model effects for anticipatory responses per trial are found in Figure 3. There were generally more anticipatory responses made during the reversal phase [main effect of Phase: χ^2^_(1, N=106)_ = 102.584, *p* = .000; Table 2]. In addition, more anticipatory responses were made on the side that was rewarded during discrimination, regardless of current phase [χ^2^_(1, N=106)_ = 23.541, *p* < .001]. We did not detect a main effect of sex, nor any higher-level interactions involving sex (all χ^2^ < 4.037, all p > .05; Table 2), There was a main effect of genotype for anticipatory responses [χ^2^_(2, N=106)_ = 6.667, *p* = .036; Fig. 2]; however, the direction of the effect was that *Syn3* deletion reduced premature responding. Homozygous wildtype mice made more anticipatory responses than the heterozygous mice (post hoc *p* = .024), and although the Cumming estimation plot comparing genotypes (Fig 2, bottom panel) suggests a mean difference separation between the homozygous null and homozygous wildtype groups, our *post hoc* analysis evaluating this difference in our model did not reach to the level of significance (*p* = .098). We believe this is a consequence of adjustments made to the estimated marginal means (EMMs) used for *post hoc* analyses within the model. EMMs are adjusted to correct for the influence of other factors in the model in order to specifically evaluate the variance explained by the factor being tested.

**Figure 2.**
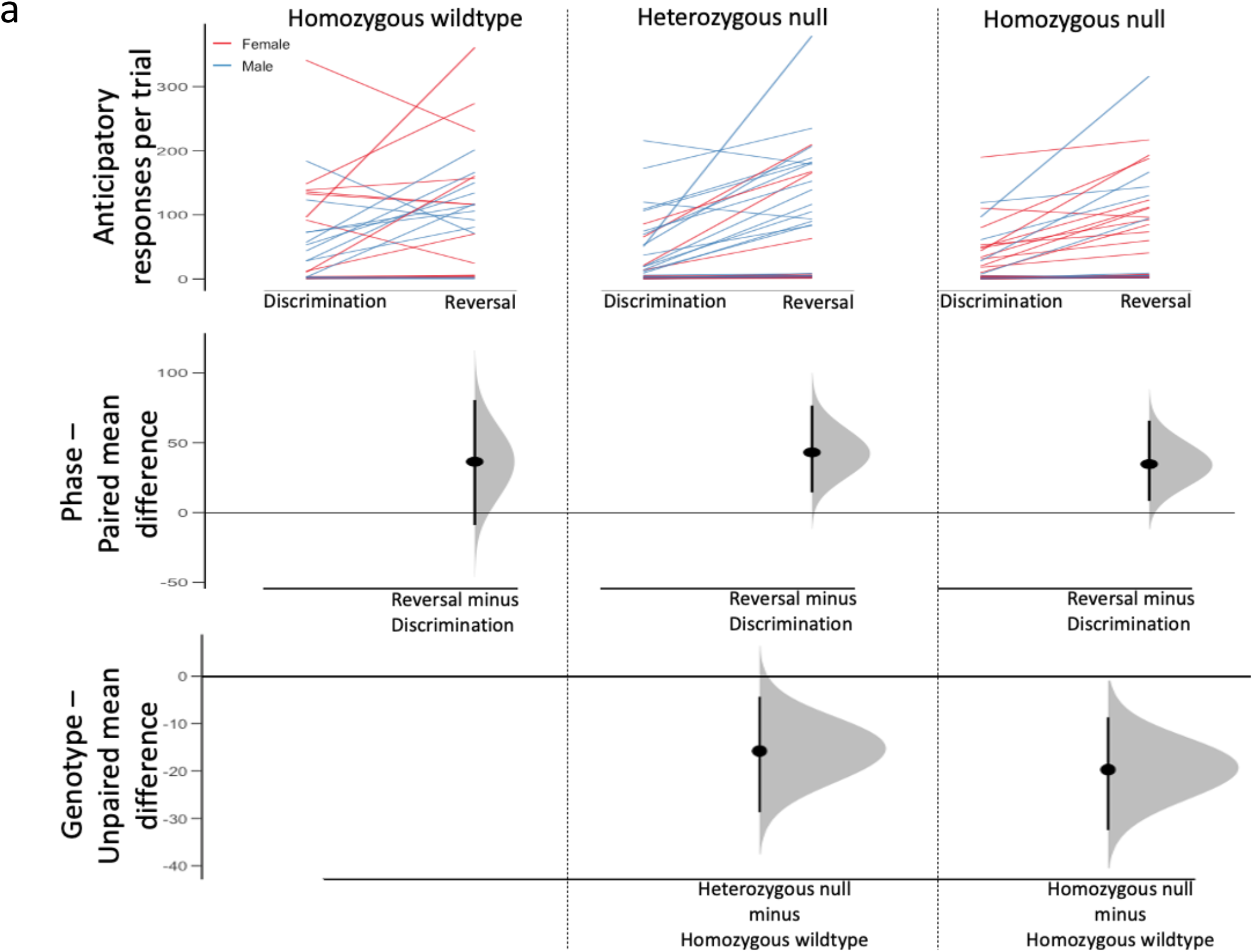
Contingency reversal increases anticipatory responses, but *Syn3* ablation reduces anticipatory responses compare to wildtype. **(a)** Cumming estimation plot displaying anticipatory responses per trial in discrimination and reversal phases within each genotype group. The line plot in the top panel shows individual data for mice in each genotype group. Paired mean differences comparing discrimination and reversal performance within each group are in the middle panel. Unpaired mean differences comparing each mutant group to the wildtype control group are shown in the bottom panel.

**Figure 3.**
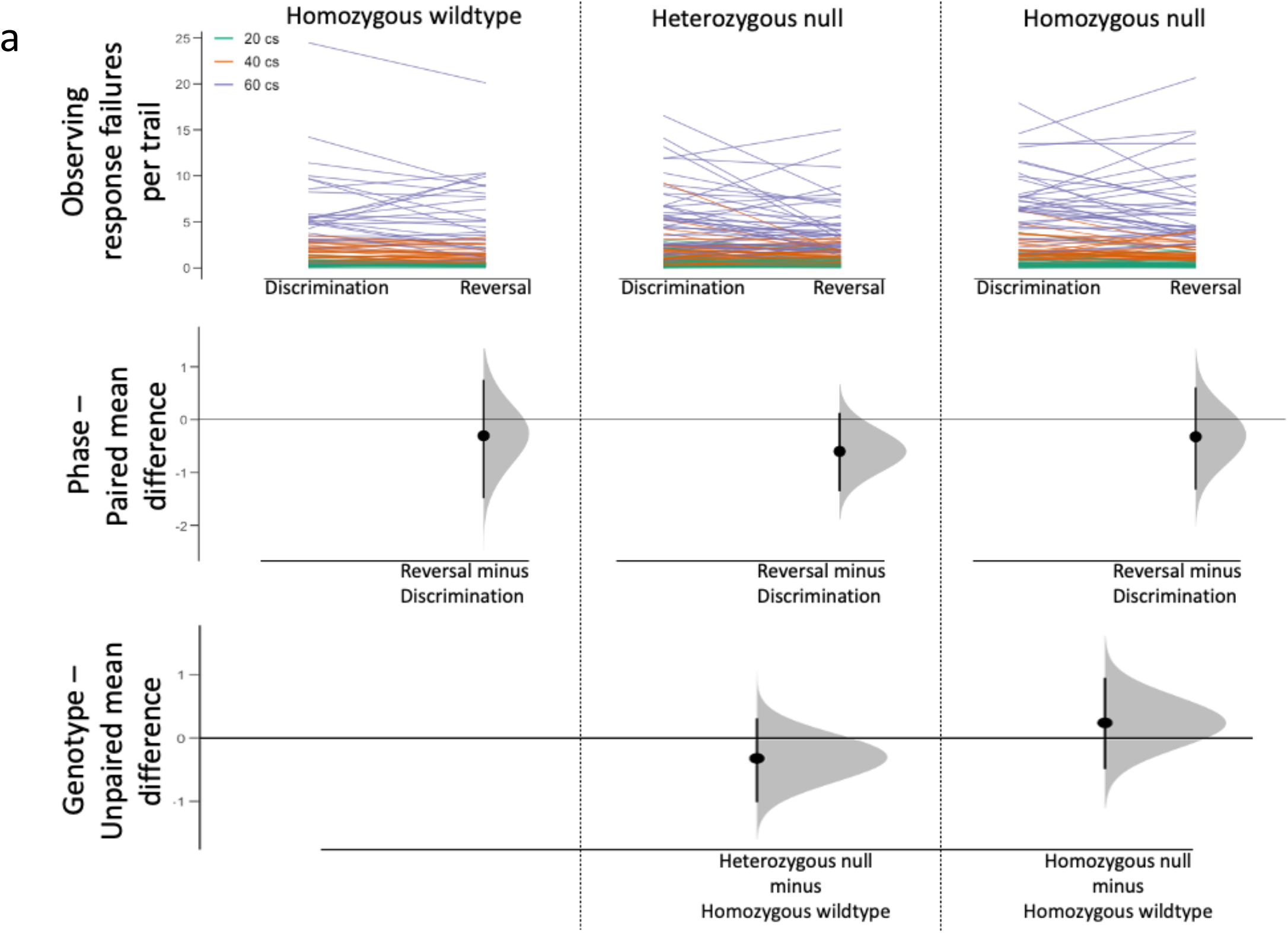
Observing response failures per trial show a probable difference between heterozygous null and homozygous null groups. **(a)** Cumming estimation plot displaying observing response failures per trial in discrimination and reversal phases within each genotype group. The line plot in the top panel shows individual data for mice each genotype group. Paired mean differences comparing discrimination and reversal performance within each group are in the middle panel. Unpaired mean differences comparing each mutant group to the wildtype control group are shown in the bottom panel.

To further investigate this issue, nested mean tables were generated to visualize the pattern of descriptive means as a function of each factor level. Descriptive means at the sex*genotype level showed the same rank-order patterns identified in our *post hoc* and estimation analyses. Specifically, females displayed the same pattern of EMMs (homozygous wildtype > homozygous null > heterozygous null), while males matched the pattern seen in genotype-based descriptive means (homozygous wildtype > heterozygous null > homozygous null). We next examined Genotype*Sex EMMs and found the same sex*genotype pattern. EMMs for genotype were then generated excluding sex from the model and the pattern was found to match that of the descriptive means, suggesting that the inclusion of sex in the model accounted for the observed effects.

### Omissions

We found a main effect of Phase [χ^2^_(1, N=106)_ = 34.308, *p* = .000; Table 2]; mice omitted fewer trials in reversal than in discrimination. No other differences were observed for trial omissions.

### OR failures

There was a main effect of Phase, such that more observing response failures occurred during the acquisition stage, than during reversal [χ^2^_(1, N=106)_ = 13.536, *p* < .001; Table 2]. Longer observing response requirements resulted in more failures per trial [χ^2^_(2, N=106)_ = 754.522, *p* < .001; Fig 3, top panel], and *post hoc* assessment revealed significant differences between all hold requirements (all pairwise *p* < .001). Genotype was also found to significantly impact observing response failures [χ^2^_(2, N=106)_ = 6.086, *p* = .048]; however, *post hoc* analyses comparing each mutant type to the wildtype control did not reveal pairwise differences. The Cumming estimation plot comparing genotypes (Fig. 3, bottom panel) shows that heterozygous null and homozygous null groups deviate from the homozygous wildtype group in opposite directions, suggesting the main effect of genotype identified in the GEE model is a difference between heterozygous and homozygous null groups.

### Latency measures

Latency model effects are reported in Table 2. Neither trial initiation latency or pellet retrieval latency was found to vary as a function of phase, genotype, sex, or their interactions (all *p* > .05).

Response latencies were longer in acquisition, than in reversal [χ^2^_(1, N=106)_ = 224.982, *p* < .001]. There was also a significant Phase*Genotype interaction [χ^2^_(2, N=106)_ = 7.254, *p* = .027]. All mice reduced their response latencies between the acquisition and reversal phases, but the effect was most pronounced in the heterozygous group. There were no differences in latency as a function of correct or incorrect responses (no effects of Side, all *p* > .05; Table 2).

### Behavioral flexibility score

Contingency reversal impacted feedback integration in a valence-dependent manner [Phase*Feedback valence interaction: χ^2^_(1, N=106)_ = 162.468, *p* < .001 Table 2]. In discrimination acquisition, mice exhibited stable behavior following positive feedback (mean = -.50, SE = .023) and flexible behavior following negative feedback (mean = .28, SE = .037), but this separation was largely lost in the reversal phase (mean positive = -.20 SE = .021; mean negative = -.19, SE = .023). A Phase*Genotype interaction was identified [χ^2^_(2, N=106)_ = 7.930, *p* = .019], and pairwise *post hoc* analysis revealed a more negative flexibility score for null mice in reversal compared to their flexibility in discrimination (*p* = .034; Fig 4a), suggesting the behavior of *Syn3* null mice were less flexible in reversal than in acquisition. We also identified Feedback valence*Sex [χ^2^_(1, N=106)_ = 9.039, *p* = .003] and Phase*Feedback valence*Sex [χ^2^_(1, N=106)_ = 6.379, *p* = .012] interaction effects. Male mice were less likely to change responses after positive feedback and more likely to update behavior after negative feedback compared to females (i.e., less flexibility after positive feedback and more flexibility after negative feedback; Fig. 4b), but as with the Phase*Feedback interaction, this difference only occurred in the discrimination acquisition phase, and differences were not apparent between groups in reversal.

**Figure 4.**
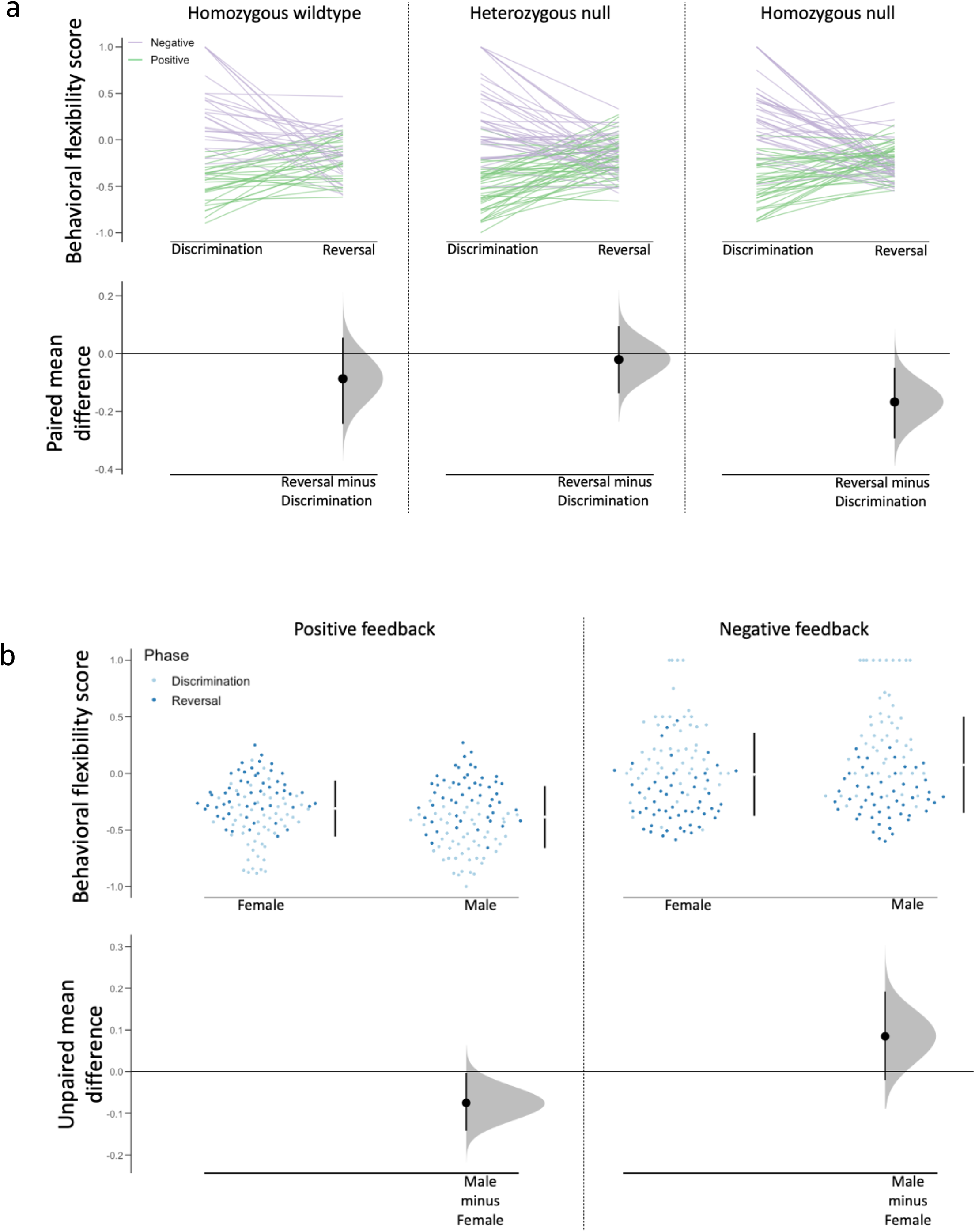
*Syn3* ablation reduces flexible behavior during contingency reversal. **(a)** Cumming estimation plot displaying behavioral flexibility during discrimination and reversal within each genotype group. The line plot in the top panel shows individual data points for mice in each genotype group. Paired mean differences comparing discrimination and reversal performance within each group. **(b)** Cumming estimation plot displaying behavioral flexibility discrimination and reversal in both sexes. The swarm plot in the top panel shows individual data points for each group with mean (±SD) displayed in the bar to the right. Unpaired mean differences comparing discrimination and reversal performance within each group.

## Discussion

In this experiment, we tested the influence of the *Syn3* gene on reversal learning performance in C57BL/6N mice using a global knockout strategy; based upon our systems genetics analysis of this trait in the BXD population (Laughlin et al., 2011), we hypothesized that mice lacking *Syn3* would display deficits in reversal learning performance compared to wildtype mice. Portions of our data support that hypothesis, but the precise effect of synapsin III deletion on reversal learning is nuanced and worthy of further exploration.

We found some evidence that reversal learning ability varies as a function of genotype. When a fold-change variable that isolates reversal-phase-specific aspects of performance was calculated, significant differences were apparent between the genotypes. Specifically, mice carrying one or two null alleles experienced a larger proportional increase in trials to criterion in reversal compared to wildtype mice. Furthermore, *Syn3* null mice exhibited less flexible behavior in reversal compared to the initial discrimination (as shown by the flexibility score analysis), again suggesting a reversal-specific deficit. These findings are consistent with our observations in BXD mice (Laughlin et al., 2011); in that study, we found that the trials to criteria under reversal testing was associated with the chromosome 10 QTL. When that measure was regressed on trials to criteria for the acquisition phase at the individual subject level, and the mean residual scores per strain (the portion of variance in the reversal data that could not be accounted for by variance in the acquisition data) were submitted to a second genome scan, we found statistically identical results, showing that the association was specific to some reversal-specific feature of behavior.

The rate of learning under the reversal condition was not the only phenotype affected by *Syn3* deletion. Wildtype mice made more anticipatory responses than either mutant group. This measure is conceptually similar to the premature responding phenotype measured in the 5-choice serial reaction time task (5CSRTT), often interpreted as an indicator of waiting impulsivity (Dalley & Ersche, 2019). At a minimum, the dissociation between behavioral flexibility assessed in reversal (impaired) and premature responding (reduced) in *Syn3* null mice suggest that these two traits are influenced by separate genetic architectures, as existing evidence already suggested (Nautiyal et al 2017). That hypothesis is further supported by our recent observation that measures of behavioral flexibility and premature responding are not genetically correlated in the Collaborative Cross recombinant inbred panel and their inbred founder strains (Bailey et al., 2021). Moreover, *Syn3* negatively affected the ability to maintain the variable duration OR. How, and/or whether, this phenotype is related to their reduced premature responding is unclear. It is also worth noting that premature responding in the 5CSRTT is typically penalized by a timeout period that lengthens the time before the start of the next trial, so inhibition of premature responses is required for optimal task performance. That is not the case in our reversal learning task; premature responses are recorded but have no programmed consequences.

In our procedure, premature responses could occur during the intertrial interval (before the OR becomes available) or during the trial initiation period (after the OR is available but before the OR duration criteria is met). Responses occurring during the trial initiation period could signal a failure to attend to the discriminative stimulus indicating a reward is now available for the correct choice (i.e., the OR aperture light turns off and the flanking lights turn on). Responding during the intertrial interval is somewhat more difficult to interpret, partially because we are unsure if these responses occurred following an attempted OR or are independent of an OR. Observing response failures, on the other hand, occurred at very low levels when the OR requirement was low, indicating the average response duration was likely higher than the minimum OR requirement. As the duration criteria increase, however, failures occur more often. Perfect success could be generated by sustaining the aperture response until the aperture light turns off on all trials, but clearly this is not the strategy used by the mice. Instead, they sustain the OR for some variable period of time that produces almost certain success at the shortest OR duration, and much lower success rates with more sustained response requirements.

Elaborating the microstructure of premature response patterns in reversal learning would lend clarity to the relation between the observing response and the premature or anticipatory response in preparations like ours that require a sequence of behavioral responses to receive a reward. If an OR failure is followed by another OR attempt, the subject is likely still attending to the relevant aperture or otherwise noted that the discriminative stimulus did not shift to the choice aperture. If an OR failure is followed by one or more anticipatory responses, the subject has shifted behaviors without attending to the lack of shift in discriminative stimuli. Though both patterns could be classified inefficient, they denote different underlying behavioral strategies. The ability to sustain a response for a sufficient duration measures a different dimension of inhibitory control than the ability to wait for choice conditions to be met before making a response. It is possible that the variable OR duration presented a greater challenge for *Syn3* null mice by introducing uncertainty into the behavioral response itself. This is functionally distinct from responding in the choice apertures prior to a choice being available, in which *Syn3* ablation seems to confer an advantage.

### Role for DA

Given the known regulatory role of *Syn3* on DA transmission (Kile et al., 2010), we can cautiously link our observed results to altered DA dynamics. Laughlin et al (2011) found Syn3 expression in the neocortex, hippocampus, and striatum was correlated with reversal learning performance (low Syn3 expression was associated with poor reversal learning performance). Synapsin III negatively regulates DA release by controlling the transfer of synaptic vesicles from the reserve pool to the ready-releasable pool (Feng et al., 2002), and mice lacking *Syn3* show enhanced striatal phasic DA release compared to wildtypes (Kile et al., 2010). Presumably, *Syn3* deficiency would also induce enhanced release in hippocampus and neocortex, but these effects have not been explicitly tested to our knowledge. Inducible and region-specific knockout strategies could be leveraged to better understand the relative contribution of corticolimbic and corticostriatal circuits to reversal learning.

DA has a complex and nuanced relationship with many behavioral phenotypes, and behavioral flexibility is no different. As noted above, inactivation of the D1-mediated direct pathway of the basal ganglia interfered with the acquisition of novel and reversed contingencies, while inactivation of the D2-mediated indirect pathway interfered selectively with reversal performance by increasing perseverative errors (Yawata et al. 2012). In a study examining DA in a compulsivity relevant behavioral phenotype, Barker et al. (2014) found that inhibiting D1 or activating D2 in the infralimbic cortex promotes behavioral flexibility.D2 agonism in the nucleus accumbens impairs flexibility (Haulk & Floresco, 2009), as does systemic antagonism of D2/D3 receptors (Lee et al., 2007). Human reversal learning performance was also found to correlate with activation of an OFC-amygdala pathway mediated by D2 receptors (van der Schaaf et al., 2013). These data indicate that there are region- and task-specific implications for DA in behavioral flexibility.

Phasic DA release has been proposed to serve as a teaching signal, encoding a prediction-error signal whereby unexpected outcomes generate an increase in phasic DA (Schultz, 2019). Steinberg et al (2013) demonstrated that optogenetic stimulation of VTA DA neurons concurrent with reward delivery produced long-lasting enhancement of cue-induced reward-seeking. Saunders et al. (2018) replicated and extended these findings, demonstrating that VTA, but not SNc DA release paired with cue presentation evoked CS approach behavior. Further, pairing optogenetic DA stimulation with reward in a behavioral economic procedure shifts the demand curve rightward and upward, indicating higher subjective value to the reward and enhanced motivation to obtain it (Schelp et al., 2017). Synapsin III negatively regulates dopamine release by regulating the transfer of synaptic vesicles from the reserve pool to the ready-releasable pool (Feng et al., 2002), and its absence enhances phasic dopaminergic tone (Kile et al., 2010). Thus, mice carrying one or more null *Syn3* alleles may experience the initial acquisition of a discrimination differently, and our test may simply not be sensitive enough to detect any reinforcement learning phenotypes during the initial acquisition stage.

### Relevance to Substance Use Disorder

Simulant drugs of abuse are widely known to act by enhancing dopaminergic activity, either by blocking DAT (e.g., cocaine) or by promoting the reverse transport of DA (e.g., amphetamine and methamphetamine). A recent systematic review and meta-analysis of the literature regarding dopaminergic alterations in stimulant users found several noteworthy changes in DA dynamics, including reduced overall DA release, reduced DAT availability, reduced D2/3 receptor availability, and possibly reduced DA synthesis (Ashok et al., 2017). Notably, striatal D2/3 receptor availability has been negatively correlated with reversal learning performance (Laughlin et al., 2011; Groman et al., 2011; Dalley et al., 2007). Theoretically, *Syn3* deletion, by amplifying phasic DA release, may lead to compensatory decreases in D2-like receptors in forebrain regions, thereby conferring a less flexible phenotype.

### Conclusions

Here we have demonstrated a role for *Syn3*, encoding synapsin III, in behavioral flexibility. C67BL/6N mice lacking functional *Syn3* alleles experienced a greater proportional cost of contingency reversal than wildtype mice and engaged in less flexible responding during the reversal phase but made fewer anticipatory responses; however, it is important to note that the reported pattern of effects may well be dependent upon the genetic background studied (Sittig et al., 2016). This suggests that *Syn3* homozygous null mice were less adaptable to changes in contingency, but this effect was independent of their waiting impulsivity phenotype.

## Acknowledgements

These studies were supported, in part, by Public Health Service grants R21-DA038377 (JDJ), T32-AA025606 (JDJ and AM) and P50-DA039841 (JDJ). The authors have no conflicts of interest to declare.

